# Regulatory architecture of gene expression variation in the threespine stickleback, *Gasterosteus aculeatus*

**DOI:** 10.1101/055061

**Authors:** Victoria L. Pritchard, Heidi M. Viitaniemi, R.J. Scott McCairns, Juha Merilä, Mikko Nikinmaa, Craig R. Primmer, Erica H. Leder

## Abstract

Much adaptive evolutionary change is underlain by mutational variation in regions of the genome that regulate gene expression rather than in the coding regions of the genes themselves. An understanding of the role of gene expression variation in facilitating local adaptation will be aided by an understanding of underlying regulatory networks. Here, we characterize the genetic architecture of gene expression variation in the threespine stickleback (*Gasterosteus aculeatus*), an important model in the study of adaptive evolution. We collected transcriptomic and genomic data from 60 half-sib families using an expression microarray and genotyping-by-sequencing, and located QTL underlying the variation in gene expression (eQTL) in liver tissue using an interval mapping approach. We identified eQTL for several thousand expression traits. Expression was influenced by polymorphism in both *cis* and *trans* regulatory regions. *Trans* eQTL clustered into hotspots. We did not identify master transcriptional regulators in hotspot locations: rather, the presence of hotspots may be driven by complex interactions between multiple transcription factors. One observed hotspot co-located with a QTL recently found to underlie salinity tolerance in the threespine stickleback. However, most other observed hotspots did not co-locate with regions of the genome known to be involved in adaptive divergence between marine and freshwater habitats.

## Introduction

It is now known that much adaptive evolution is underlain by changes in regions of the genome regulating gene expression, rather than in the protein coding regions of the genes themselves (Pavey *et al.* 2010). Recent work has demonstrated that much variation in gene expression is heritable, and thus evolvable via selection (e.g. Ayroles *et al*. 2009, Powell *et al.* 2013, Leder *et al.* 2015). Correspondingly, studies using model species have found that the genetic polymorphisms underlying phenotypic variation are typically not within genes (Flint and Mackay 2009). Variation in gene expression has been shown to underlie several well-documented cases of phenotypic and/or adaptive divergence. These include plumage coloration and beak shape in birds (Mallarino *et al.* 2011; Poelstra *et al.* 2015), mimetic wing patterns in butterflies (Reed *et al.* 2011; Hines *et al.* 2012), and flower colour (Durbin *et al.* 2003). Further, differences in gene expression patterns have been found to correlate with adaptive divergence in multiple species (e.g. Bernatchez *et al.* 2010; Barreto *et al.* 2011). Dysregulation of gene expression due to interactions amongst regulatory loci has potential to cause reduced fitness of inter-population hybrids and thus contribute to reproductive isolation (Ellison and Burton 2008; Turner *et al.* 2014). However, it may also promote hybrid speciation by enabling hybrids to exploit new niches (Lai *et al.* 2006).

The genetic architecture of gene expression regulation can be investigated by treating expression variation as a quantitative trait and identifying the genomic locations associated with it (termed ‘expression quantitative trait loci’ or ‘eQTL’). Such studies have shown that the expression of a gene can be regulated by multiple genomic regions, which are traditionally classified as either *cis* or *trans*. *Cis* regulators, including promoters that activate transcription and enhancers that influence transcription levels, are located close to the regulated gene(s). They contain binding sites for regulatory molecules (proteins or mRNA) that are produced by more distant, *trans*, regulators. As *cis* regulators are expected to affect only one or a few focal genes, while *trans* regulators may have pleiotropic effects on many genes, *cis* and *trans* regulators are subject to different evolutionary dynamics. *Cis* regulatory changes are expected to be important drivers of local adaptation (Steige *et al.* 2015), while *trans* regulatory variation is considered more likely to be under purifying selection (Schaefke *et al.* 2013 but see also Landry *et al.* 2005 for discussion of *cis-trans* coevolution). Correspondingly, *trans* regulatory polymorphisms tend to affect gene expression less strongly than *cis* polymorphisms, and their effects are more likely to be non-additive (Zhang *et al.* 2011; Gruber *et al.* 2012; Schaefke *et al.* 2013; Meiklejohn *et al.* 2014; Metzger *et al.* 2016). Nevertheless, work in multiple species has demonstrated an important role for both *cis* and *trans* polymorphism in shaping expression variation (Cubillos *et al.* 2012; Meiklejohn *et al.* 2014; Guerrero *et al*. 2016) and the role of *trans* variation may have been underestimated due to the higher statistical power required to detect it (Mackay *et al.* 2009; Clément-Ziza *et al.* 2014). Interactions involving *trans* regulators may be particularly important in reducing the fitness of inter-population hybrids (Turner *et al.* 2014). Supporting the pleiotropic role of *trans* regulators, a ubiquitous feature of eQTL studies is the identification of ‘*trans* eQTL hotspots’, genomic locations associated with expression variation in many distant genes which are thought to harbour one or more important *trans* regulators (Wu *et al.* 2008; Clément-Ziza *et al.* 2014; Meiklejohn *et al.* 2014).

The threespine stickleback (*Gasterosteus aculeatus*) is an important model in the study of adaptive evolution. Ancestral anadromous populations of threespine stickleback have repeatedly and independently colonized freshwater throughout the Northern Hemisphere (Taylor and McPhail 2000; Mäkinen *et al.* 2006). Sympatric and parapatric freshwater populations may exploit different habitats (Schluter and McPhail 1992; Roesti *et al.* 2012). The species is also distributed throughout semi-marine environments with large temperature and salinity gradients, such as estuaries and the brackish water Baltic Sea (McCairns and Bernatchez 2010; Guo *et al.* 2015; Konijnendijk *et al.* 2015). Successful colonization of these diverse habitats necessitates behavioural, morphological and physiological adaptation to novel environmental conditions including changed temperature, salinity, oxygen, light, parasite and predator regimens, a process that can occur rapidly (Kitano *et al.* 2010; Barrett *et al.* 2011; Terekhanova *et al.* 2014; Lescak *et al.* 2015; Huang *et al.* 2016, Rennison *et al.* 2016). Parallel adaptations between independently founded freshwater populations frequently involve the same regions of the genome and arise from pre-existing genetic variation in the marine population (Colosimo *et al.* 2005; Hohenlohe *et al.* 2010; Jones *et al.* 2012; Liu *et al.* 2014; Conte *et al.* 2015, but see DeFaveri *et al.* 2011; Leinonen *et al.* 2012; Ellis *et al.* 2015; Ferchaud and Hansen 2016). Local adaptation in environmentally heterogeneous habitats such as the Baltic Sea (Guo *et al.* 2015) and lake-stream complexes (Roesti *et al.* 2015) has been shown to involve the same genomic regions. Evidence suggests that much of this adaptation may be due to changes in gene regulation rather than protein structure (Jones *et al.* 2012). In addition, plasticity in gene expression in response to different environmental conditions may facilitate initial colonization of novel habitats.(McCairns and Bernatchez 2010; Morris *et al.* 2014). Leder *et al.* (2015) recently demonstrated substantial heritability of expression variation, over thousands of genes, within a Baltic Sea threespine stickleback population, confirming that it can be shaped by local selection. One well-documented locally adaptive trait, reduction of the pelvic girdle, is known to be underlain by variation in the *cis* regulatory region of the *Pitx1* gene (Chan *et al.* 2010), and *cis*-regulatory variation at the *Bmp6* gene underlies divergent tooth number between a freshwater and marine population (Cleves *et al.* 2014). Differences in levels of thyroid hormone between freshwater and marine sticklebacks, which are connected to different metabolic rates between the two environments, are associated with *cis* regulatory variation at the *TSH*β*2* gene (Kitano *et al.* 2010). Recently, Di Poi et al (2016) showed that differences in behaviour and response to stress between marine and freshwater sticklebacks may be modulated by variation in the expression of hormone receptors. Otherwise, the architecture of gene expression regulation in the threespine stickleback and its role in adaptive evolution is only starting to be explored (Chaturvedi *et al.* 2014).

Understanding the regulatory pathways underlying variation in gene expression, and how this gene expression variation influences the phenotype, will improve our understanding of how organisms can adapt to novel environments and, thus how adaptive diversity is generated. In the stickleback, for example, it is unknown whether regulatory loci involved in local adaptation are clustered on the regions of the genome that show repeated divergence in independent marine-freshwater colonizations. Here, we perform the first genome-wide study of this regulatory architecture in the threespine stickleback, by mapping QTL underlying the variation in expression of several thousand genes in a population from the Baltic Sea. We examine transcription in the liver, a metabolically active tissue that expresses many genes potentially involved in physiological adaptation to different aquatic habitats.

## Methods

### Experimental crosses

We used a multi-family paternal half-sib crossing design for QTL mapping. Crossing procedures have previously been detailed in Leinonen *et al.* (2011) and Leder *et al.* (2015). In short, 30 mature males and 60 gravid females were collected from the Baltic Sea for use as parents. Each male was artificially crossed with two females, producing 30 half-sib blocks each containing two full-sib families. Families were reared in separate 10L tanks with density standardized to 15 individuals per tank, temperature at 17 ± 1°C and 12:12h light/dark photoperiod. At the age of six months, ten offspring from each family (5 treated, 5 controls) were subject to a temperature treatment as part of a related experiment (control: constant 17°C; treatment: water gradually heated from 17°C to 23°C over 6 hours, see Leder *et al.* 2015), and immediately euthanized for DNA and RNA collection.

### RNA preparation, microarray design, and data normalization

RNA preparation, gene expression microarrays, hybridization, and normalization procedures are described in detail in Leder *et al.* (2009, 2015). Briefly, total RNA was isolated from offspring liver tissue using standard protocols. RNA that passed quality thresholds was labelled (Cy3 or Cy5) using the Agilent QuickAmp Kit, with equal numbers of individuals within family groups (control & temperature-treated; males & females) assigned to each dye. Labelled RNA was hybridized to a custom 8x15K microarray, with sample order randomized (Agilent Hi-RPM kit). Labelling, hybridization, and scanning was performed at the University Health Network in Toronto, Canada. Images of the arrays were acquired, image analysis was performed, and array quality was assessed as detailed in Leder *et al.* (2015). Post-processed signals were standardized across arrays using a supervised normalization approach, implemented in the package 'snm' for R/Bioconductor (Mecham *et al.* 2010; R Core Team 2015). Dye, array and batch (i.e. slide) were defined as ‘adjustment variables’; sex, family and temperature treatment were defined as ‘biological variables’. Following normalization, individual intensity values more than two standard deviations from their family-by-treatment mean, and probes with missing values for an entire family or >10% of individuals were removed. The final dataset contained 10,527 expression traits (10,495 genes plus 32 additional splice variants) and 563 individuals (158 control females; 125 control males; 152 treated females; 128 treated males).

### Genotyping-by-Sequencing

For genotyping-by-sequencing of parents (n = 90) and offspring (n = 580) we used the method of Elshire *et al.* (2011) with an additional gel excision step to improve size selection. DNA was extracted from ethanol preserved fin tissue (parents) or frozen liver tissue (offspring) and DNA concentrations were measured using a NanoDrop ND-1000 spectrophotometer. DNA (80 ng) was digested with the restriction enzyme Pst1 1.5 U (New England Biolabs) and 1x NEB buffer 3, 1x bovine serum albumin (BSA) and dH_2_O (3.3 µl) in a thermocycler (37°C, 2h; 75°C, 15min; 4°C, 10min). The digested DNA was ligated to adapters with T4-ligase 0.6x (New England Biolabs), 1x Ligase Buffer, 21 µl dH_2_O and 50 nM of pooled forward and reverse adapters, which were prepared according to Elshire *et al.* (2011; ligation program: 22°C, 1h; 65°C, 30min; 4°C, 10min). Up to one hundred and four unique barcodes were used in each library to label individual samples. The ligation products were pooled into libraries and purified with a QIAquick PCR Purification Kit (Qiagen). The purified libraries were PCR amplified with the following components: Purified ligated library (20µl), reaction buffer 1x, MgCl_2_ 1.5nM (Bioline), primer mix 0.5 µM, dNTPs (Fermentas) 0.4μM, BioTaq 0.05 U (Bioline) and dH_2_O (20µl) (Amplification program: [72°C, 5min; 4 cycles [95°C, 30s; 95°C, 10s; 65°C, 30s; 70°C, 30s]; 11 cycles [95°C, 10s; 65°C, 30s; 72°C, 20s]; 72°C, 5min; 4°C, 10min). Lastly, we performed a manual size selection by loading 40 µl of the amplified library on a gel (MetaPhor [Lonza] 2.5 %, 150 ml, 100 V. 1.5 h) and cutting the 300-400 bp range from the resultant smear. The DNA was extracted from the gel with a QIAquick Gel Extraction Kit (Qiagen). The cleaned product was again separated on a gel, cut and cleaned.

All products were sequenced with paired-end reading on the Illumina HiSeq2000 platform. Six hundred and fifty individuals, multiplexed into ten separate libraries (maximum library size = 104 individuals), were sequenced at the Beijing Genomics Institute (BGI); 55 individuals (including duplicates) were sequenced at the Finnish Institute for Molecular Medicine or at the University of Oslo.

### Variant calling

Reads were split by barcode, and barcodes removed, using a custom perl script. Low quality bases were removed from the reads via window adaptive trimming using Trim.pl (available: http://wiki.bioinformatics.ucdavis.edu/index.php/Trim.pl, Illumina quality score ≤ 20). Paired-end reads for each of these individuals were aligned to the BROAD S1 stickleback genome using BWA aln/sampe (v 0.6.2) with default parameters (Li and Durbin 2009). The threespine stickleback genome comprises 21 assembled chromosomes plus 1,823 un-placed genomic scaffolds. Unmapped reads, and reads with non-unique optimal alignments, pair-rescued alignments, or any alternative suboptimal alignments, were discarded from resulting SAM files. SAM files were converted to sorted BAM files using samtools 0.1.18 (Li *et al.* 2009) and variants were called within each paternal family using the samtools mpileup function with extended BAQ computation (options: -AED, max-depth 500), in combination with bcftools (Li *et al.* 2009). We did not degrade mapping quality for reads with large numbers of mismatches as we found this to reject high-quality reads due to fixed polymorphisms between our European stickleback samples and the North American stickleback genome. Indel and multi-allelic variants were discarded. Initial filters based on SNP quality and variability within and across families resulted in list of 26,290 candidate bi-allelic SNPs for further analysis. Samtools and bcftools, applied to each paternal family separately, were then used to call each individual for the genotype at each of the 26,290 sites. Sites at which bcftools identified multiple variant types (SNPs, indels and multi-base polymorphisms) within and among families were removed, leaving 25,668 successfully genotyped variant sites.

### Genotype quality control

Vcftools (Danecek *et al.* 2011) was used to recode genotypes with a genotype quality phred score (GQ) < 25 or a sequencing depth (DP) < 8 or > 1000 to missing. Vcf files for all families were merged and the merged file converted to the input format for Plink 1.07 (Purcell *et al.* 2007). For SNPs on all autosomal chromosomes and the pseudoautosomal region of Chromosome 19 (see below), the following filters were applied in Plink: hwe (based on founders only) < 0.01, maximum missing genotypes = 0.25, minor allele frequency > 0.05, offspring with > 70% missing data removed. Adjacent SNPs in complete linkage disequilibrium were manually consolidated into a single locus, with combined SNP information used to call genotypes.

Several approaches were used check for sample contamination or errors in barcode splitting and family assignment: in Plink, the *mendel* option was used to screen families for Mendelian errors, and sample relatedness was examined by graphically visualizing genome-wide IBD-sharing coefficients generated by *genome*; the program SNPPIT (Anderson 2012) was used to assign individuals to parents, based on five independent datasets of 100 SNPs; and 220 SNPs on Stratum II of Chromosome 19 (see below) were examined for their expected pattern in males and females (all heterozygous in males vs. all homozygous in females).

The stickleback Chromosome 19 is a proto-sex chromosome (Peichel *et al.* 2004; Roesti *et al.* 2013; Schultheiß *et al.* 2015), with a normally recombining pseudo-autosomal domain (approximately 0-2.5mB), a non-recombining domain in the male version (Stratum I, approximately 2.5-12mB) and a domain largely absent in the male version (Stratum II, approximately 12-20mB). For Stratum I, parental and offspring genotypes were inspected manually in order to identify the male-specific allele and this was recoded to a unique allele code (‘9’) for the purposes of linkage map construction. Where the male-specific allele could not be identified, all genotypes within a family were re-coded as missing. Genotypes were also inspected manually for Stratum II, and any SNP found to be heterozygous in males was excluded. All remaining Stratum II SNPs were considered to be hemizygous in males, and one of the alleles was also recoded as ‘9’.

### Linkage map construction

We constructed a linkage map using the improved version of Crimap (Green *et al.* 1990, available: http://www.animalgenome.org/tools/share/crimap/). Remaining Mendelian errors in the dataset were removed using the *set-me-missing* option in Plink. For each SNP, the number of informative meiosis was examined using Crimap, and markers with < 150 informative meioses or within 500bp of one another were discarded.

The initial map build included 6,448 markers. Where applicable, SNPs were ordered according to the modified genome build of Roesti *et al.* (2013). We attempted to position all previously un-placed scaffolds containing at least two genotyped SNPs on to the map. Scaffolds were assigned to chromosome on the basis of LOD score using the Crimap function *two-point*, and then positioned using a combination of information from pilot Crimap *builds, chrompic*, and *fixed* together with known start and end points of previously assembled scaffolds (Roesti *et al.* 2013). Information from *chrompic* and *fixed* were also used to confirm the orientation of scaffolds newly placed by Roesti *et al.* (2013). Once all possible scaffolds had been placed, recombination distance between ordered SNPs was estimated using *fixed.* To refine the map, we iteratively removed SNP genotypes contributing to implied double crossovers within a 10 cM interval (presumed to be genotyping errors), and SNPs generating recombination distances of >1cM per 10,000 bp and recalculated distances using *fixed.* Remaining regions of unusually high recombination on the map were investigated by examining whether removal of individual SNPs altered map distance.

### eQTL identification

Expression QTL (eQTL) were identified using an interval mapping approach (Knott *et al.* 1996) implemented in QTLMap 0.9. 0 (http://www.inra.fr/qtlmap; QTLMap option: – *data-transcriptomic*). Offspring with missing genotypes at > 60% of the markers in the linkage map were removed from the analysis. We applied linkage analysis assuming a Gaussian trait distribution (QTLMap option: – *calcul = 3*), and included dye, temperature treatment, and sex as fixed factors in the model. Due to the relatively small size of some of our half-sib families, we examined sire effects only, with a separate QTL effect estimated for each sire. Excluding dam effects is expected to reduce our power of eQTL detection, as fewer parents will be segregating for each QTL.

A fast algorithm was used to identify phase and estimate transmission probabilities at each chromosomal location (Elsen *et al.* 1999, QTLMap option: *–snp*). Autosomes and the pseudoautosomal portion of the sex chromosome were scanned at 1cM intervals, and the presence of QTL on a chromosome was assessed using a likelihood ratio test (LRT) under the hypothesis of one versus no QTL. Chromosome-wide LRT significance thresholds for each trait were identified empirically, by permuting fixed effects and traits amongst individuals within families and recalculating LRT scores (5000 permutations). As the combination of 5000 permutations x 10,332 traits x 21 chromosomes was computationally prohibitive, we first performed permutations on a subset of 200 expression traits to establish a LRT threshold below which identified QTL were unlikely to be significant at chromosome-wide p < 0.05 (LRT = 55), and then used permutations to assess significance of all QTL above this threshold. The non-pseudo-autosomal region of the female Chromosome 19 can be considered analogous to the X chromosome; identification of QTL in this region requires estimation of dam effects and was therefore not performed. The 95% confidence interval for each QTL was estimated using the drop-off method implemented in QTLMap 0.9.7, which returns flanking map positions plus their nearest marker.

### Cis vs. trans eQTL

To discriminate *cis* vs. *trans* QTL, we compared inferred QTL location to the position of the expressed gene according to the BROAD *G. aculeatus* genome annotation v. 1.77 (available: http://ftp.ensembl.org/pub/release-77/gtf/gasterosteus_aculeatus/). All positions on the BROAD annotation were re-coded to positions on our modified chromosome assemblies. For genes on scaffolds un-anchored to our assembly, we also used information on chromosomal scaffold locations available in the recently published map of Glazer *et al.* (2015). Any eQTL on a different chromosome from the regulated gene was considered *trans*. For eQTL on the same chromosome as the gene, we initially considered two alternative threshold distances for an eQTL to be considered *trans* (> 1Mb following Grundberg *et al.* 2012, or > 10Mb following van Nas *et al*. 2010). For the 1Mb threshold, we observed strong enrichment of significant *trans* eQTL on the same chromosome as the regulated gene, indicating that these were actually mis-identified *cis* eQTL; we therefore selected the conservative 10Mb threshold. In practice, examination of our results showed that 95% CI of eQTL sometimes extended further than this 10Mb threshold. Considering median 95% CI (approximately 1Mb), we therefore classified a QTL as *trans* if the SNP closest to the upper or lower 95% confidence bounds of that QTL was further than 9.5Mb from the regulated gene. Following Johnsson *et al.* (2015) we applied a local significance threshold (chromosome-wide p < 0.01) for evaluation of possible *cis*-QTL and a genome-wide significance threshold (genome-wide p < 0.021, = chromosome-wide threshold of 0.001 * 21 chromosomes) for evaluation of possible *trans*-QTL. Although this significance threshold is permissive, we considered it acceptable as our aim was to analyse the eQTL distribution across the genome rather than to identify individual QTL-locus associations. Similar significance thresholds have been used for eQTL detection in comparable studies (e.g. Whiteley *et al.* 2008).

To ask whether the effect of variation in *trans* regulatory sites was more often non-additive than the effect of variation in *cis* regulatory sites, we examined the narrow sense heritability (*h*^*2*^) and dominance proportion of genetic variance (*d*^*2*^) estimated for each expression trait by Leder *et al.* (2015) and provided in the Supplementary Data for that paper.

### Genes with plastic vs. non-plastic expression

To investigate whether genes exhibiting an alteration in expression level in response to a temperature stress treatment (i.e. those exhibiting environmental plasticity) had a different underlying regulatory architecture to those not exhibiting such a response, we divided genes into a ‘responding’ and ‘non-responding’ group based on the results provided in the Supplementary Data for Leder *et al.* (2015) and compared the frequency and position of *cis* and *trans* eQTL between the two groups.

### Evaluation of eQTL hotspots

As all identified eQTL had wide 95% confidence intervals, meaning that physically close eQTL positions could be due to the effect of the same locus (see below), we evaluated potential eQTL hotspots by counting eQTL within 5cM bins across the genome (‘hotspot size’ = number of eQTL). Where the number of 1cM bins within a chromosome was not a simple multiple of 5, bin sizes at the start and/or end of the chromosome were increased to 6 or 7. To obtain an empirical significance threshold above which clusters of eQTL could be considered a ‘hotspot’, we simulated the expected neutral distribution of eQTL across the genome using a custom script. We performed 5000 simulations: for each, we assigned *n* eQTL (where *n* = relevant number of significant eQTL) randomly across the 3,062 1cM bins of the genome and then summed them into 5cM (or larger) bins as described above. Conservatively, we compared the size of hotspots in the real data to the size distribution of the largest hotspot observed over each of the 5000 simulations.

### Association of eQTL with regions under selection

Hohenlohe *et al.* (2010), Jones *et al.* (2012), and Terekhanova *et al.* (2014) documented parallel regions of the genome divergent between marine and freshwater sticklebacks on Chromosomes 1, 4 (three regions), 7, 11 and 21. We investigated whether these regions harboured important *trans* regulators that might contribute to adaptation to different aquatic habitats by comparing the location of these regions with the location of our identified *trans* eQTL hotspots. We also compared hotspot location to regions of the genome inferred by Guo *et al.* (2015) to be involved in adaptive differentiation amongst different stickleback populations in the Baltic Sea.

### Ortholog identification

In order to maximize the functional information available, we identified human orthologues for *G. aculeatus* genes. As a first attempt, we used BioMart (Durinck *et al.* 2005; Smedley *et al.* 2009) to identify human orthologues and obtain the HGNC symbols for the human genes. When BioMart failed to return a human orthologue, protein BLAST searches were used to identify orthologues using the Ensembl human protein database. The identifier conversion tool, db2db, from bioDBnet (https://biodbnet-abcc.ncifcrf.gov/db/db2db.php) was used to convert between Ensembl identifiers and HGNC gene symbols when needed (Mudunuri *et al*. 2009).

### Hotspot annotation

To identify regulatory genes physically associated with an eQTL hotspot, we defined hotspot confidence boundaries as being the most frequently observed 95% confidence limits of all significant eQTL centred in the hotspot. We used AmiGO2 (Carbon et al. 2009) to identify ‘molecular function’ or ‘biological process’ Gene Ontology (GO) terms associated with transcriptional regulation by applying the search term [transcription regulation and – pathway]. We then used BioMart to examine all genes within the hotspot boundaries for any of these GO annotations, using the HGNC symbols as input. As an important transcriptional regulator generating a hotspot might itself be regulated by the hotspot rather than physically present within it, we repeated this analysis for all genes with eQTL mapped to the hotspot (*cis* eQTL significant at chromosome-wide p < 0.01; *trans* eQTL significant at genome wide p < 0.021). We used DAVID (Huang *et al.* 2009a; b).to examine GO term enrichment for the sets of genes with *trans* QTL mapping to each hotspot, using the 9,071 genes on the microarray with identified human orthologues as the background. To increase our sample size, we lowered our stringency and examined all genes with *trans* eQTL mapping to the hotspot locations at genome-wide p < 0.057 (chromosome-wide p < 0.0027).

### Upstream regulator and functional interaction analyses

To search for regulatory genes which may be responsible for the expression variation in genes with identified *trans* eQTL, we used the upstream regulator analysis in the Ingenuity Pathway Analysis (IPA) software (Qiagen). This analysis uses a Fisher’s Exact Test to determine whether genes in a test dataset are enriched for known targets of a specific transcription factor. We used the human HGNC symbols as identifiers in IPA. First we examined all genes that had a significant *trans*- eQTL mapping to any location at a genome-wide p < 0.021 (chromosome –wide p < 0.001). To investigate in more detail the upstream regulators potentially involved in generating eQTL hotspots, we lowered our stringency and also examined all genes with *trans* eQTL mapping to the hotspot locations at genome-wide p < 0.057 (chromosome-wide p < 0.0027).

Since transcription is typically initiated by a complex of genes rather than a single transcription factor, we examined functional relationships among the identified upstream regulators for each hotspot (Table S7b), the genes located within a hotspot, and the genes with significant eQTL mapping to that hotspot (Table S3; *cis* eQTL significant at chromosome-wide p < 0.01, *trans* eQTL significant at genome-wide p < 0.021), using STRING v10 (Jensen *et al.* 2009, http://string-db.org/). We searched for evidence of functional relationships from experiments, databases and gene co-expression, and applied a minimum required interaction score of 0.4.

## Results

### Genotyping by sequencing

Sufficient numbers of reads were obtained for 620 of the 670 individuals sent for sequencing. Fifteen of these individuals failed initial quality control steps. For the 605 sticklebacks (88 parents, 517 offspring) that were retained for analysis, we obtained a total of 583,032,024 raw paired reads (40,357 – 11,940,726 per individual, median=834,286). Approximately 67% of these reads remained aligned to the stickleback genome following removal of reads with non-unique optimal alignments, any alternative suboptimal alignments, or pair-rescued alignments (range 36.2% - 78.8%, median = 70.1%). Raw read and alignment statistics for each individual are provided in Table S0.

### Linkage map construction

Following SNP calling and quality control steps 13,809 of the original 25,668 SNPs, genotyped in 605 individuals (mean number of offspring per family = 18), were available for linkage map construction. Following removal of markers with < 150 informative meioses or within 500bp, 6,448 SNPS were included in the initial map build. The final sex-averaged linkage map spanned 3,110 cM Kosambi (including the complete Chromosome 19) and included 5,975 markers, of which approximately 45% were located at the same map position as another marker (Figure 1, Figure S1, Table S1). Forty-three previously un-placed scaffolds (10.35 mB) were added to the chromosome assemblies of Roesti *et al.* (2012, Table S2). Thirty-five of these scaffolds were also recently added to the stickleback assembly in an independent study by Glazer *et al.* (2015). Although there were some differences in scaffold orientation, location of the new scaffolds was almost completely congruent between the two maps (Table S2). For QTL detection with QTLMap, the map was reduced to 3,189 SNPs with unique positions (average inter-marker distance = 0.98cM, Table S1).

**Figure 1.**
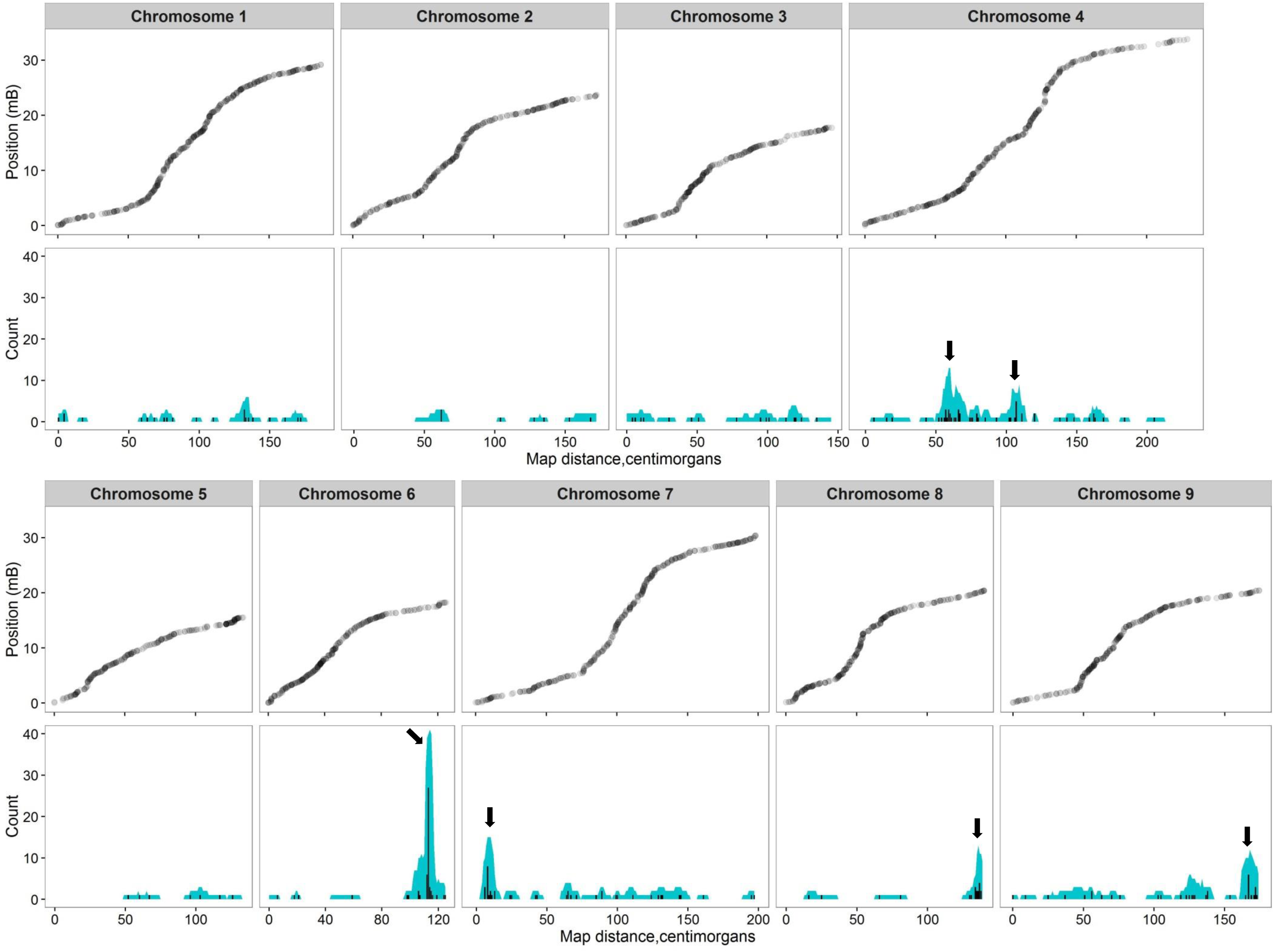
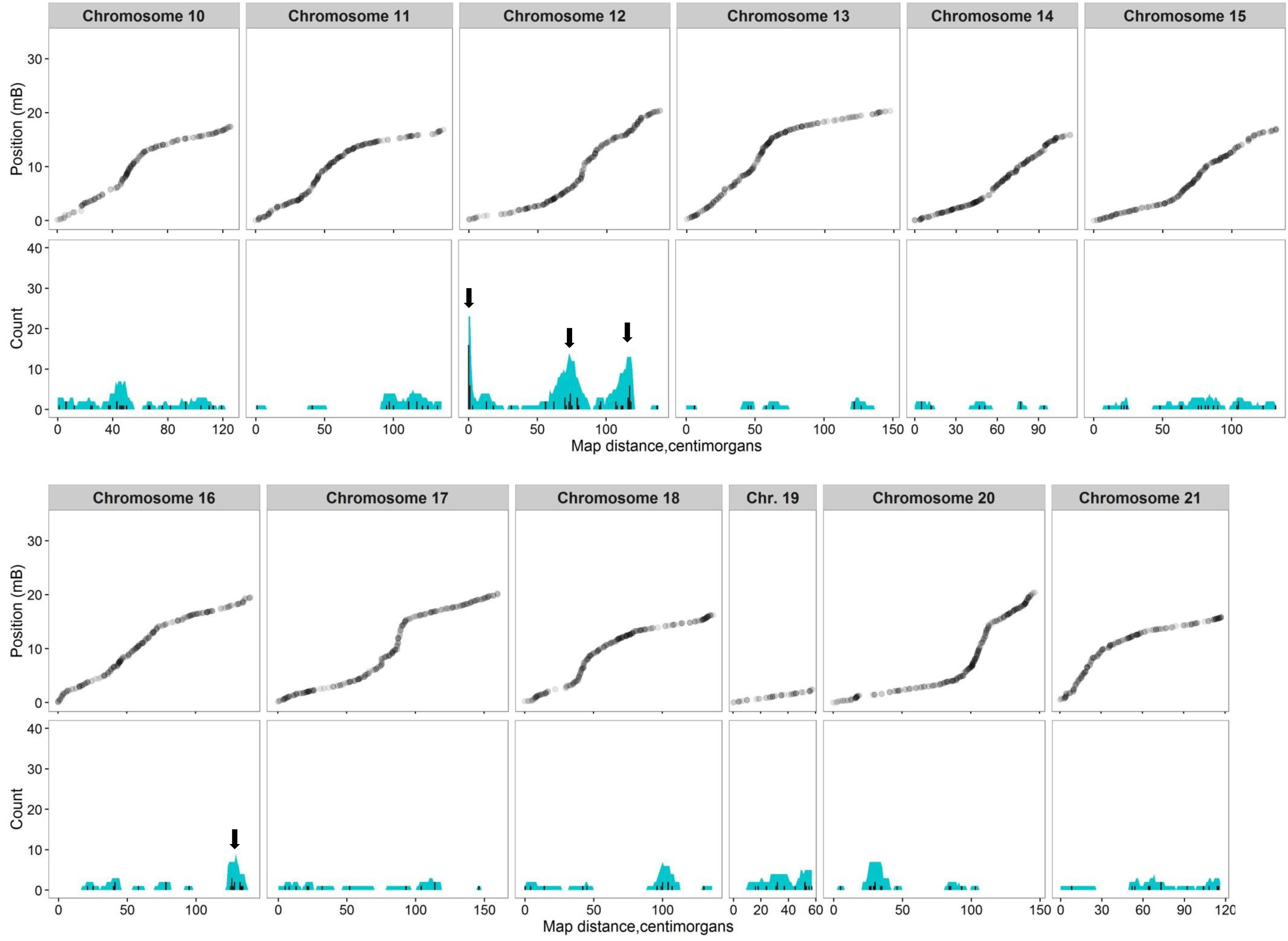
Position of SNP markers along each chromosome (top) and location of *trans* eQTL hits for all assayed genes (bottom). Black bars show the number of eQTL hits at each 1cM Kosambi interval along the chromosome. Blue shading shows the number of eQTL with 95% confidence intervals overlapping each 1cM interval. Arrows indicate the location of eight significant *trans* eQTL hotspots. Figure created using ggplot2 (Wickham 2009) in R.

### Identification of cis and trans eQTL

Expression data were available for 500 of the 517 genotyped offspring. Twenty-six of these offspring had > 60% missing genotype data and were removed from the analysis. As we found that missing values in the expression trait file caused QTLMap to over-estimate the LRT statistic, we eliminated these from the dataset by removing one additional individual and 195 expression traits. Eighty-eight genotyped parents, 473 genotyped and phenotyped offspring (mean no. offspring per family = 15.8, mean proportion of missing genotypes in offspring = 0.11; maximum = 0.56), and 10,332 expression traits were retained for the analysis. At chromosome-wide p < 0.01, we identified 5,366 eQTL associated with 4,507 expression traits (43.7% of the 10,322 expression traits examined, Table S3). Based on our recoded gene positions, we classified 2,335 of these as *cis* eQTL, 2,870 as *trans* eQTL, and 161 as unknown – that is, the expressed gene was located on a scaffold that had not been assigned to a *G. aculeatus* chromosome by either this study or Glazer *et al.* (2015; Table S3, Table S4). Four hundred and seventy four of the *trans* eQTL were significant at genome-wide p < 0.021. Of these, 84.5% mapped to a chromosome other than the one containing the regulated gene. After application of this genome-wide significance threshold for *trans* eQTL, 2,858 expression traits (27.7% of those examined) remained associated with one or more significant *cis* or *trans* eQTL. Of these, 79.4% were associated with a *cis* eQTL, 13.9% with one or more *trans* eQTL, 2.3% with both a *cis* and a *trans* eQTL and 4.4% with eQTL of unknown class (Table S3). The physical distribution across the genome of the 2,858 loci with significant *cis* or *trans* eQTL is shown in Figure S1. Mean 95% confidence interval of significant eQTL was 10.2 cM (range 1-86cM), approximating 1.77Mb (range 0.03 – 22.19Mb). Overall, *trans* regulated expression traits did not exhibit more dominance variance than *cis* regulated loci (*trans* regulated loci, mean *h*^*2*^ = 0.31, mean *d*^*2*^ = 0.16; *cis* regulated loci: mean *h*^*2*^ = 0.37, mean *d*^*2*^ = 0.18; values from Leder *et al.* 2015).

### Trans eQTL hotspots

*Trans* eQTL (significant at genome wide p < 0.021) were not evenly distributed across the genome and we identified ten 5cM bins, located on seven different chromosomes, as containing eQTL clusters (7 or more eQTL; p < 0.012 based on the largest hotspot observed in neutral simulations; Figure 1). A particularly large eQTL hotspot (36 *trans* eQTL within the 5cM bin) was identified close to one end of Chromosome 6, three hotspots (23, 7, and 8 *trans* eQTL) were present at separate locations on Chromosome 12, two hotspots (10 and 7 *trans* eQTL) were located on Chromosome 4, and remaining hotspots were located near the ends of Chromosomes 7, 8, 9 and 16 (14, 10, 8 and 7 *trans* eQTL). To eliminate the possibility that distant *cis* eQTL mis-classified as *trans* were contributing to observed hotspots, we repeated the analysis with the 401 *trans* eQTL that were on a different chromosome to their regulatory target – nine out of the ten hotspots were still present (6 or more eQTL, p < 0.038; second Chromosome 4 hotspot, with 5 eQTL, no longer significant). Physical hotspot boundaries were assigned from inspection of eQTL hits and 95% confidence intervals as follows: Chromosome 4, 55-67cM (‘Chr4a’, 4,630,680-6,394,113bp); Chromosome 4, 104-113cM (‘Chr4b’, 15,643,256-17,021,069bp), Chromosome 6, 111-116cM (‘Chr6’, 17,238,934-17,469,219bp); Chromosome 7, 5-12cM (‘Chr7’, 396,541-1,107,393bp); Chromosome 8, 134-139cM (‘Chr8’, 19,917,746-20,316,565bp); Chromosome 9, 165-174cM (‘Chr9’, 19,822,078-20,440,410bp); Chromosome 12, 0-1cM (‘Chr12a’, 0-337,849bp); Chromosome 12, 72-79cM (‘Chr12b’, 5,853,981-7,440,742bp); Chromosome 12, 109-119cM (‘Chr12c’, 15,551,555-17,229,387bp); Chromosome 16, 123-130cM (‘Chr16’, 17,658,526-18,257,571bp).

### Genes with plastic vs. non-plastic expression

Following FDR correction, 4,253 genes were found by Leder *et al.* (2015) to exhibit a significant change in expression in response to a temperature treatment. We identified significant eQTL underlying 1131 of these genes (Table S3; eQTL type: 79.8% *cis*, 12.7% *trans*, 2.6% both, 4.9% unknown). The distribution of the 177 significant *trans* eQTL across 5cM bins indicated four hotspots (5 or more eQTL, p < 0.01, Figure S2), all of which had been previously observed in the full dataset. The Chromosome 16 hotspot was greatly increased in relative importance (Chr4b: 6 eQTL; Chr 6: 12 eQTL; Chr12a: 9 eQTL; Chr16: 7 eQTL).

### Association of eQTL with regions under selection

None of our identified eQTL hotspots overlapped parallel regions of the genome divergent between marine and freshwater sticklebacks identified by Hohenlohe *et al.* (2010), Jones *et al.* (2012), and Terekhanova *et al.* (2014), or with the clusters of morphological QTL on Chromosome 20 (Miller *et al.* 2014, Table S5). However, one genomic region identified as divergent between marine and freshwater populations by Terekhanova *et al.* (2014) alone overlapped with the Chr12b eQTL hotspot. Only nine of the 297 genes inferred by Guo *et al.* (2015) as being under selection amongst Baltic Sea populations experiencing different temperature and salinity regimens overlapped observed eQTL hotspots (Chr4a, Chr4b, Chr7, Chr9 and Chr12b, Table S5).

### Hotspot annotation

We identified human orthologues for 16,315 of the 20,787 protein-coding genes annotated on the Broad stickleback genome (78.5%, Table S4). There were 393 genes with human annotation physically located within the designated boundaries of the eleven hotspots (Table S5). Of these, 70 (17.8%) had a GO term related to transcription regulation (Table 1, Table S6). In addition, 21 genes with significant *cis* eQTL or *trans* eQTL mapping to a hotspot had GO terms related to transcriptional regulation (Table 1, Table S6). Following correction for multiple testing we found no significant GO term enrichment amongst any group of genes *trans* regulated by the same eQTL hotspot.

**Table 1.**
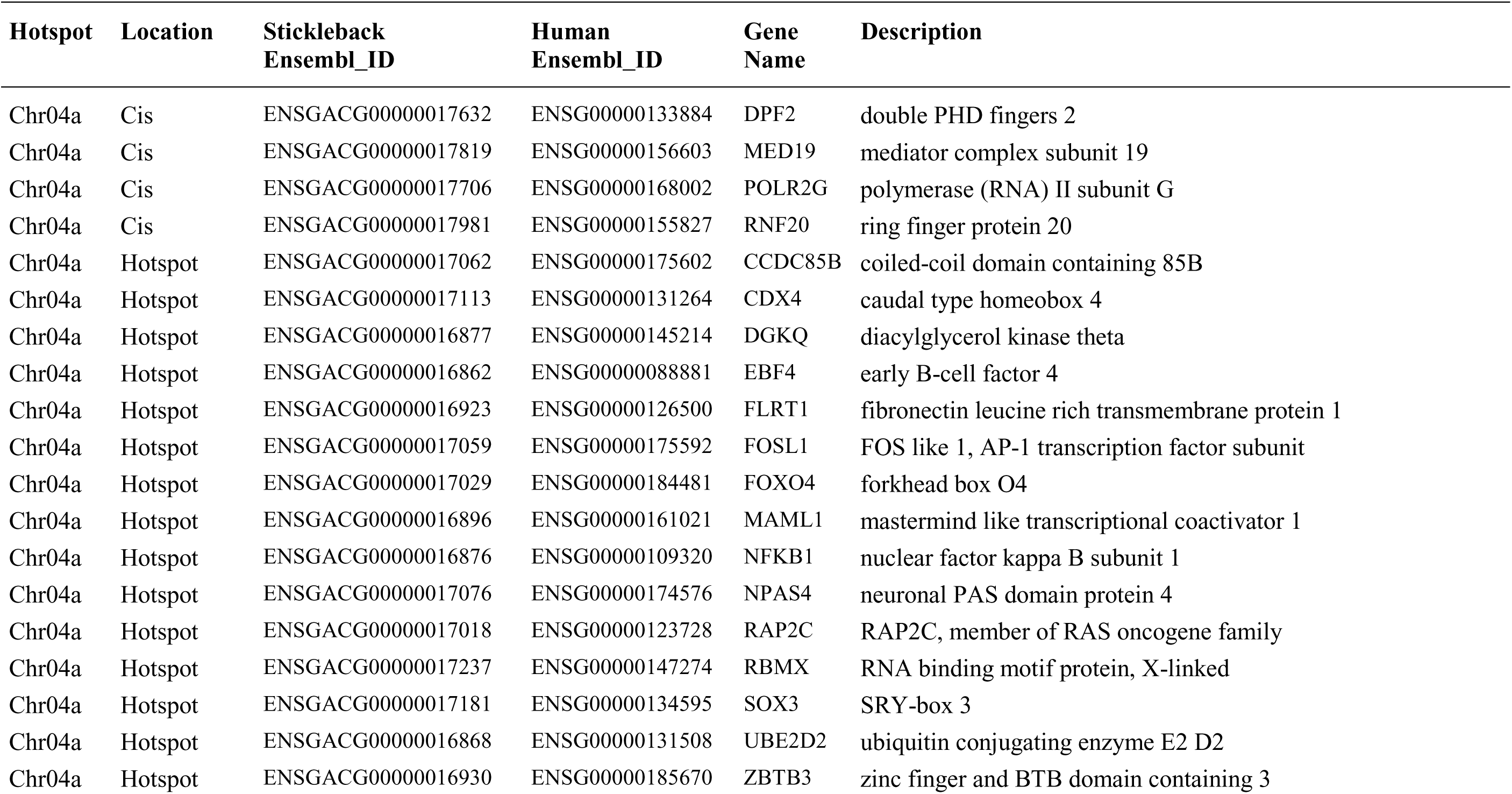
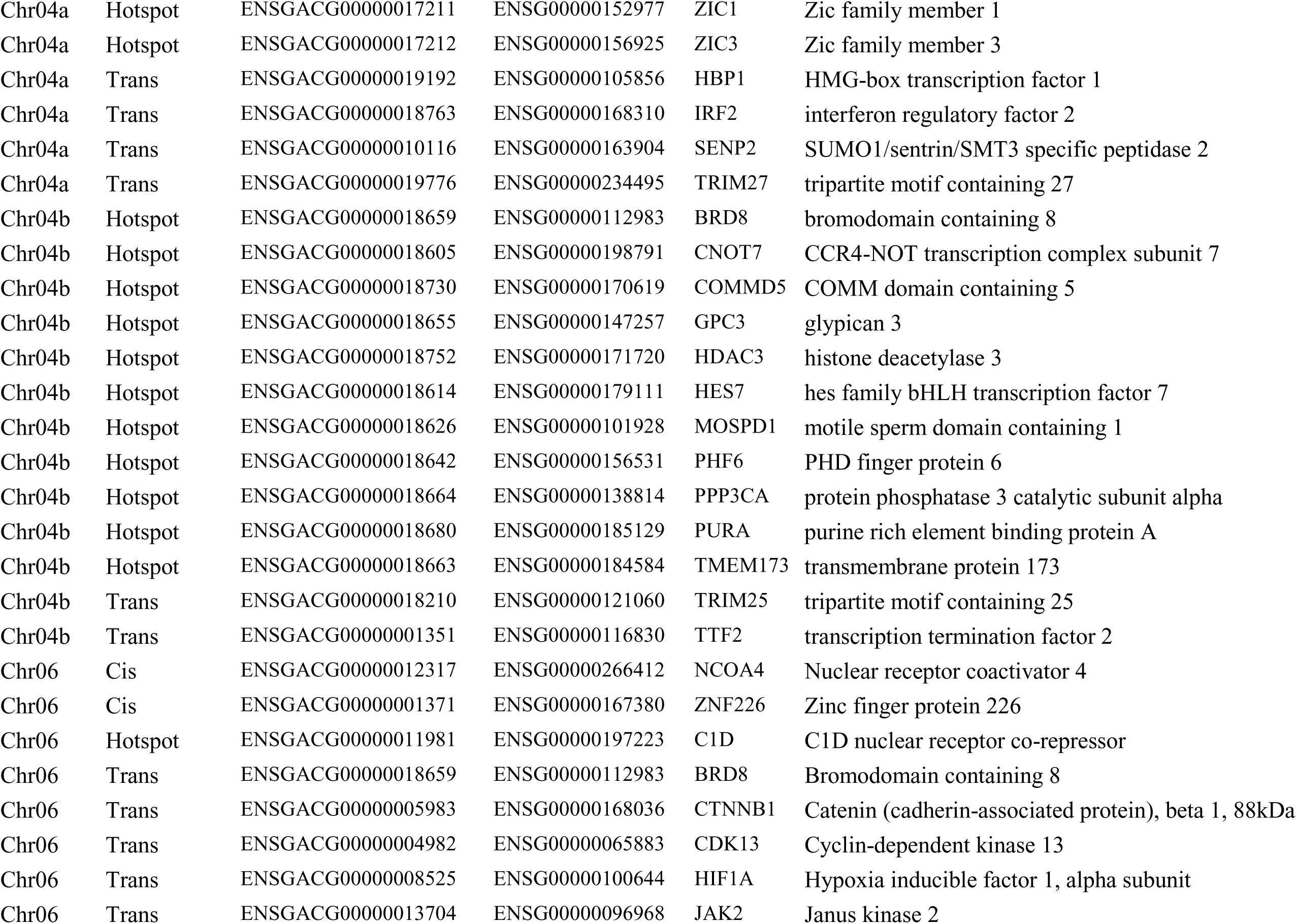
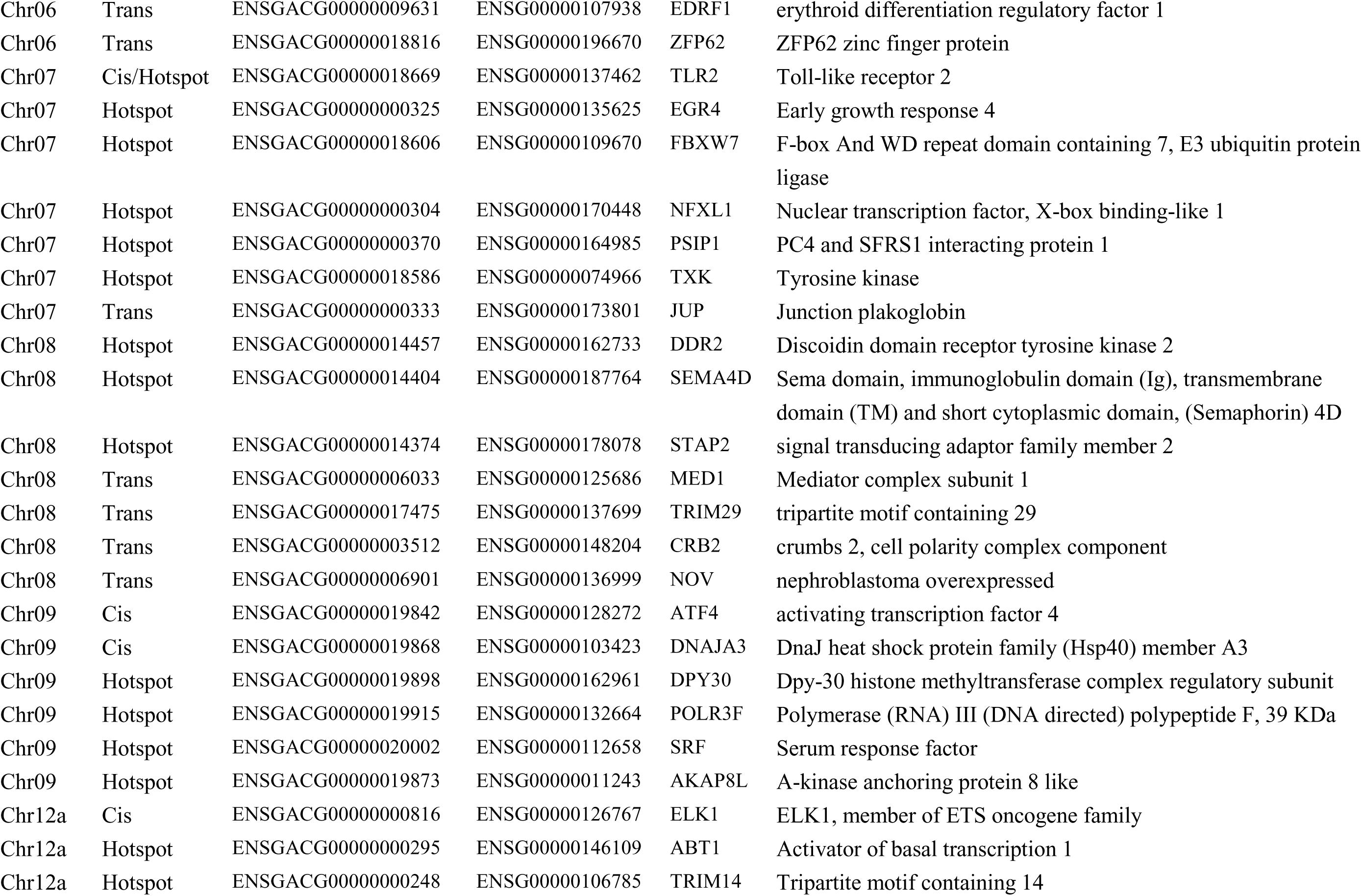
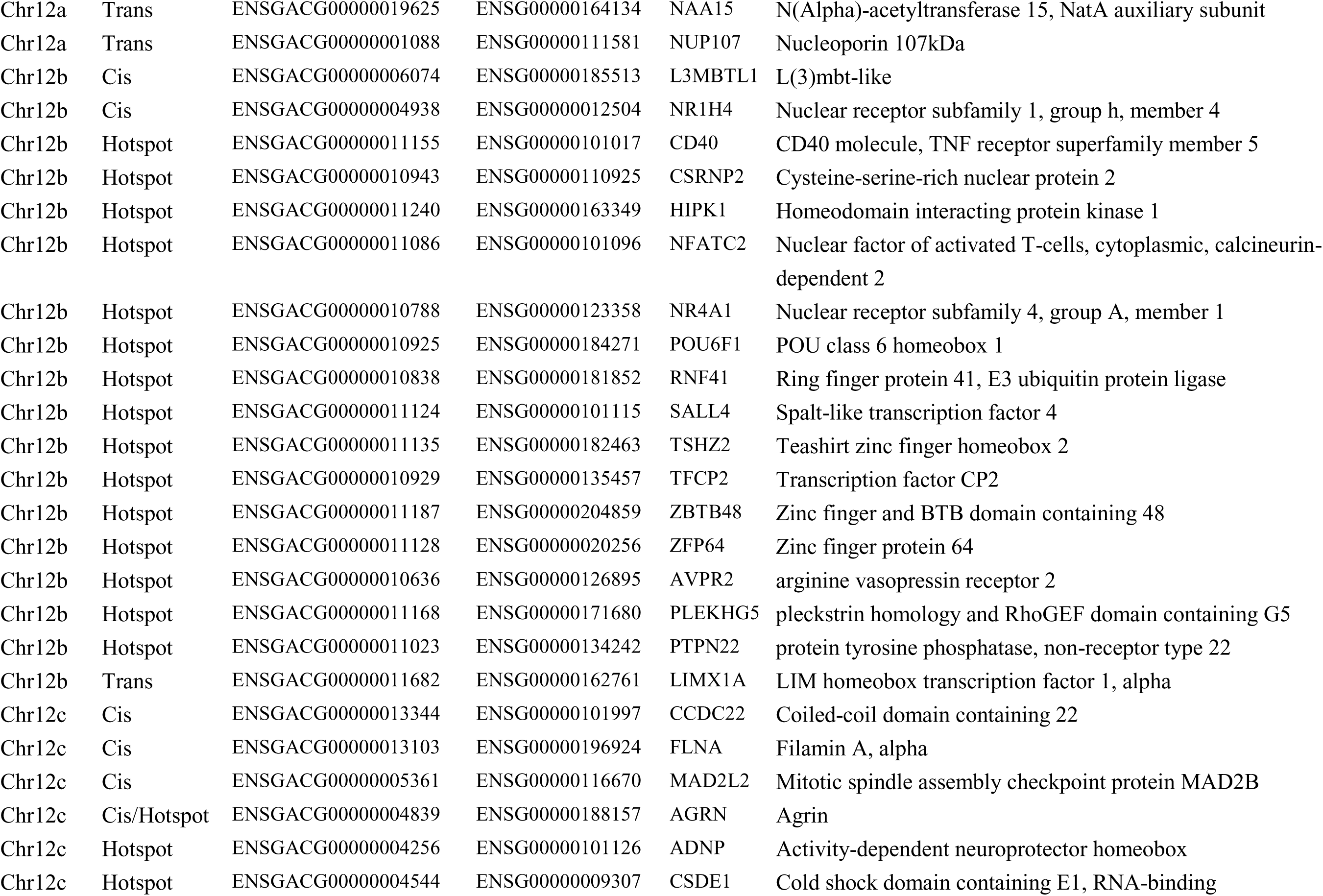
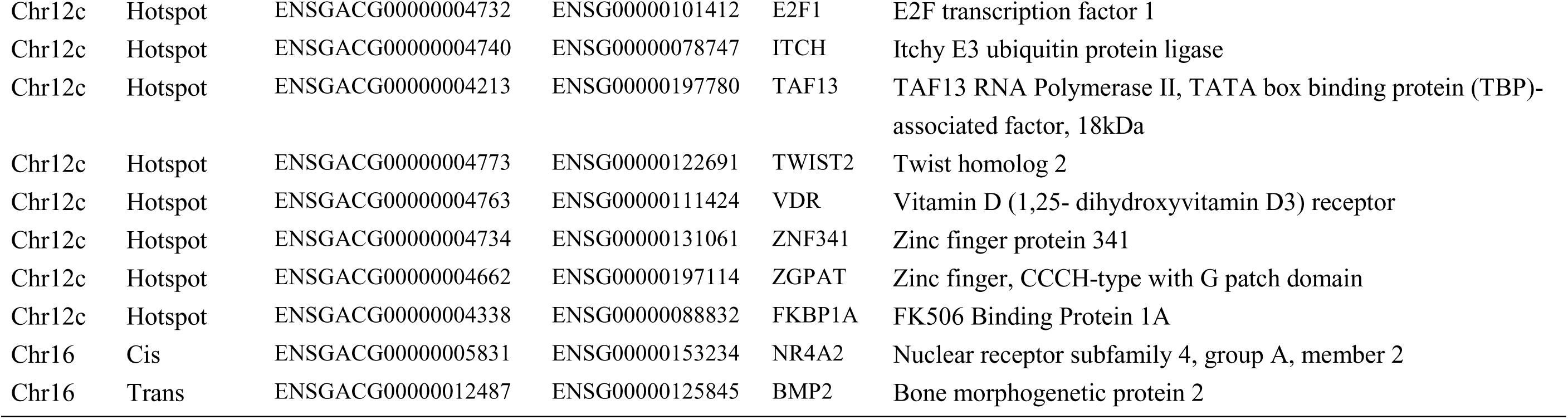
Known transcriptional regulators associated with identified eQTL hotspots. Human orthologues of stickleback genes were identified using BioMart. Location is as follows: ‘Hotspot’: annotated gene is in genomic region of hotspot; ‘*Cis* ’: gene is cis-regulated by hotspot at chromosome wide p<0.01; ‘*Trans* ’: gene is trans-regulated by hotspot at genome-wide p<0.021.

### Upstream regulator and functional interaction analyses

When examining all 405 genes with *trans* eQTL significant at genome wide p < 0.021, 79 significantly enriched upstream regulators were identified using IPA (Table S7). In total, these regulators had 208 of the genes in the dataset as known targets. Hepatocyte nuclear factor 4α (HNF4A) was identified as a particularly important regulator (p = 9.3*10^−8^), with 70 (33.7%) of these genes as downstream targets. Other highly enriched regulatory factors included one cut homeobox 1 (ONECUT1; p = 3.2*10^−5^; 16 target genes), Nuclear Receptor Subfamily 4 Group A Member 1 (NR4A1; p = 2.0*10^−4^; 11 genes), Signal Transducer And Activator Of Transcription 5B (STAT5B; p = 5.5*10^−4^; 10 genes), Krüppel-like factor 3 (KLF3; p = 8.7*10^−4^; 15 genes), estrogen receptor 1 (ESR1; p = 1.8*10^−3^; 37 genes); Hepatocyte nuclear factor 1α (HNF1A; p = 1.9*10^−3^; 17 genes); CAMP Responsive Element Binding Protein 1 (CREB1; p=3.2*10^−3^; 19 genes) and myc proto-oncogene protein (MYC; p = 3.4*10^−3^; 30 genes). The full list of 79 significant upstream regulators is in Table S7.

To identify upstream regulators that could be contributing to the ten eQTL hotspots we further examined all genes that had *trans* eQTL mapping to the hotspots at genome-wide p< 0.057 (1120 genes). One hundred and ninety-two different enriched upstream regulators were identified for these genes (Table S7b). For genes with *trans* eQTL mapping to the Chr4b, Chr6, Chr12a and Chr12c hotspots, HNF4A remained an important regulator. Only five of the identified upstream regulators were physically located within a hotspot (NFKB1, Chr4a; SOX3, Chr4a; SRF, Chr9; NFATC2, Chr12b; NR4A1, Chr12b). Five had significant *cis* or *trans* eQTL mapping to a hotspot (IRF2, Chr4a *trans*; NCOA4, Chr6*cis*; HIF1A, Chr6*trans*; JUP, Chr7*trans*; ELK1, Chr12a*cis*). None of these ten hotspot-associated regulatory proteins were identified as significant upstream regulators for the sets genes with *trans* eQTL mapping to the same hotspot – in other words, their presence did not appear to be causative of the observed hotspots.

When the enriched upstream regulators, genes with cis eQTL mapping to a hotspot at chromosome-wide p < 0.01, and genes with *trans* eQTL mapping to a hotspot at genome wide p < 0.021 were examined in STRING, multiple protein-protein interactions were found (Figure 2, Figure S4). In particular for the Chr6 hotspot we found a complex interaction network that included eight molecules *trans*-regulated by this hotspot (in order of connectivity: CTNNB1, HIF1A, CASP3, BRD8, CDK13, EIF3C, JAK2, UCK1), two molecules *cis*-regulated by the hotspot (C1D and B3GNT2), and multiple molecules inferred as important upstream regulators by IPA (Figure 2).

**Figure 2:**
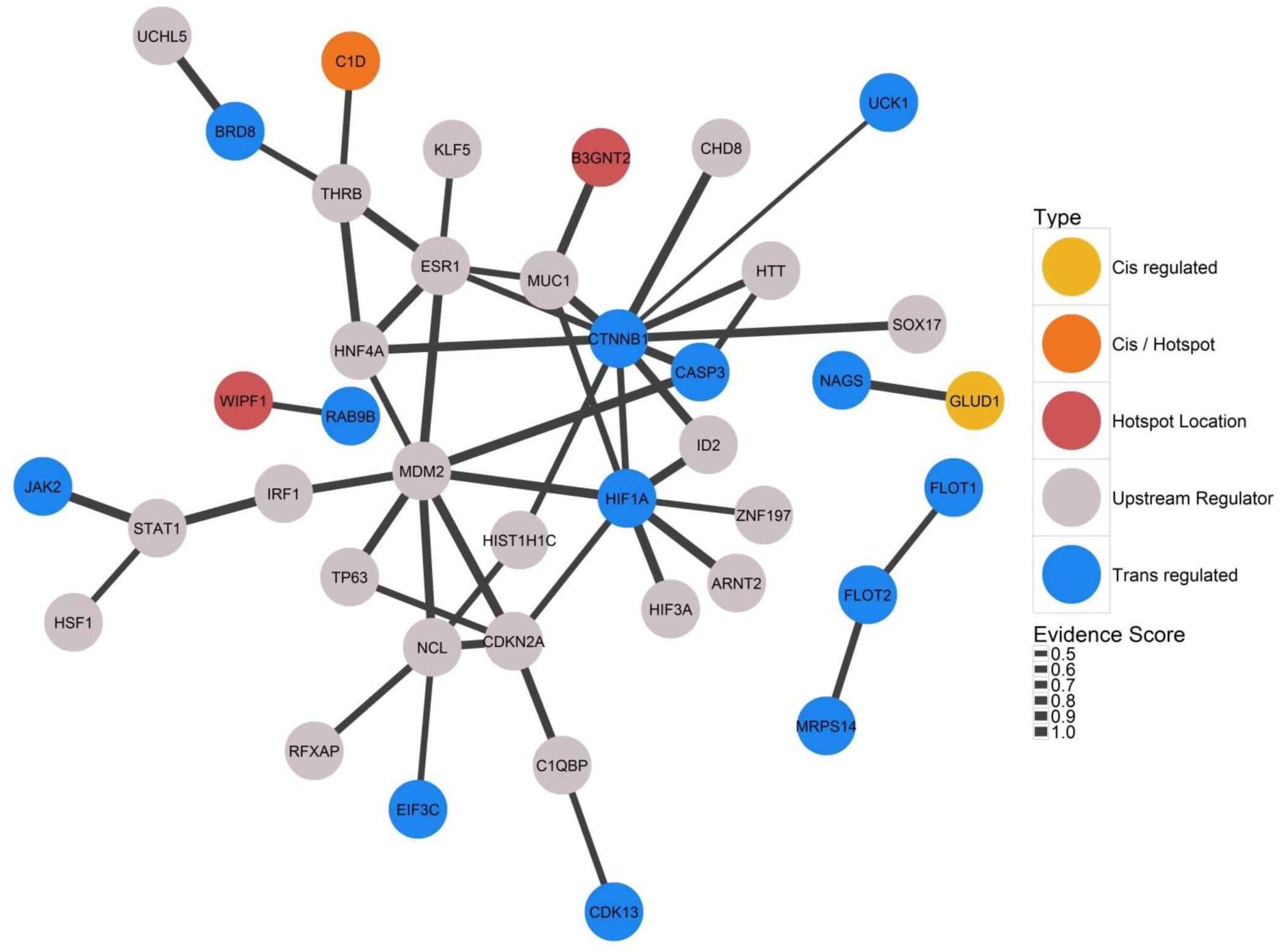
Networks of known protein-protein interactions inferred by String 10 for proteins associated with the Chr6 hotspot. ‘Upstream Regulator’: significantly enriched upstream regulator identified when examining genes *trans*-regulated by the hotspot using IPA; ‘Hotspot Location’: protein is coded by a gene physically located in the hotspot; ‘Trans regulated’: protein is *trans* regulated by an eQTL mapping to the hotspot and significant at genome-wide p<0.021; Cis/Hotspot: both present in and significantly *cis* regulated by the hotspot. Interactions not involving an identified upstream regulator are not shown.

## Discussion

In this study we identified regions of the genome underlying variation in gene expression in a population of threespine stickleback from northern Europe. We used a genotyping-by-sequencing approach to generate an improved linkage map, and applied interval mapping to identify eQTL. Our new map was independent of that recently constructed by Glazer *et al.* (2015), and the congruent placement of scaffolds between the two maps confirms the reliability of these new genome assemblies. Our map covered a substantially larger distance in cM than those of Roesti *et al.* (2013) and Glazer *et al.* (2015), probably due to differences in experimental design. Nevertheless, for our Baltic Sea population, we observe very similar patterns of recombination rate variation across and between chromosomes as found by Roesti et al. (2013) for freshwater sticklebacks from central Europe and Glazer *et al.* (2015) for marine-freshwater crosses from western North America, (Figure S1). Thus, the large scale pattern of recombination rate variation across the genome may impose, and/or be under, similar evolutionary constraints throughout the range of the species.

Using a chromosome-wide significance threshold for *cis* regulatory loci and a genome-wide threshold for *trans* loci, we identified eQTL for just over a quarter of the 10,332 expression traits examined. Because at least 74% of these expression traits exhibit significant heritable variation (Leder *et al.* 2015), and gene expression is commonly regulated by multiple eQTL, we expect that a much larger number of underlying eQTL remain undetected due to low statistical power. Despite expectations that *trans* regulatory regions might be under purifying selection due to their potentially pleiotropic effect, and that the effect of *trans* eQTL on expression will be weaker than that *cis* eQTL, we found many cases where gene expression was influenced by regulatory variation in *trans* but not in *cis*. This suggests that a frequently-used approach of detecting local selection by examining patterns of differentiation at markers linked to genes that are adaptive candidates (e.g. DeFaveri et al. 2011, Shimada et al. 2011) may fail to identify such selection as it is acting to change gene expression via *trans* regulatory regions. We did not observe any difference in additive vs dominance variance underlying genes found to be regulated in *cis* vs. those regulated in *trans*. However this may again be due to low statistical power to detect many of the underlying eQTL: genes are expected to be influenced by a large number of eQTL, meaning that the observed heritable variation is generated by a combination of additively and non-additively acting regulatory regions.

The *trans* eQTL that we detected were not randomly distributed across the genome but instead clustered into multiple eQTL hotspots. This observation is a ubiquitous feature of eQTL studies and is thought to indicate the existence of ‘master regulators’ acting in *trans* to influence many genes. However apparent eQTL hotspots may also arise as a statistical artefact as a result of many false positive QTL when testing thousands of expression traits in combination with spurious correlation between these traits due to uncorrected experimental factors (Wang *et al.* 2007; Breitling *et al.* 2008). Disentangling gene expression correlation that is due to common underlying regulatory architecture from that caused by experimental artefacts is a difficult analytical problem that we are unable to fully address here (Joo *et al.* 2014). Therefore, we caution that these hotspots should be verified using other stickleback populations and different approaches.

The parents for this study came from a genetically diverse marine population of threespine stickleback (DeFaveri *et al.* 2013). Local adaptation of threespine sticklebacks to freshwater has been demonstrated to arise, at least partly, from selection on standing genetic variation in the marine environment. Further, QTL underlying morphological divergence between marine and freshwater populations have been demonstrated to have pleiotropic effects (Rogers *et al.* 2012; Miller *et al.* 2014), and frequently co-localize with regions of the genome found to be under parallel selection amongst independent freshwater colonisations. One way in which these regions could exert such pleiotropic effects is by harbouring loci that influence the expression of many genes, i.e. eQTL hotspots. However, only one of the *trans* eQTL hotspots found in this study (Chr12a) overlapped with genomic regions repeatedly found to be associated with marine/freshwater divergence by Hohenlohe *et al.* (2010), Jones *et al.* (2012), or Terekhanova *et al.* (2014).

Several studies, however, indicate that adaptation to novel aquatic environments may also involve parts of the genome outside these large target regions (DeFaveri *et al.* 2011; Leinonen *et al.* 2012; Ellis *et al.* 2015; Ferchaud and Hansen 2016; Erickson *et al.* 2016). The QTL underlying physiological adaptations to different aquatic environments in sticklebacks have not been well characterized. Recently, Kusakabe *et al* (2016) identified a significant QTL associated with salinity tolerance (indicated by gill sodium plasma levels) on Chromosome 16, which overlaps our Chr16 *trans* eQTL hotspot. Interestingly, this also appears to overlap with a Chromosome 16 QTL underlying gill raker morphology identified by Glazer *et al* (2015). Based on transcription levels, Kusakabe *et al* (2016) identified ten candidate causal genes at the QTL location; we found *cis* regulatory variation for four of these genes (CLN5, IGFBP5, RABL3, NDUFA10) and a fifth (GDP-like) had a *trans* eQTL mapping to the Chr4a hotspot. However, Kusakabe *et al* (2016) did not investigate genes located elsewhere on the genome that may be trans-regulated by this Chromosome 16 QTL. Our results also show that all genes with *trans* eQTL mapping to the Chr16 hotspot exhibit a plastic response to the temperature treatment. Thus, the Chr 16 eQTL hotspot may be involved in physiological adjustment to several environmental variables.

Identifying eQTL directly implicated in local adaptation in sticklebacks was not our experimental aim, and it is possible that regulatory hotspots acting in tissues or life stages that we did not examine have a role in stickleback adaptive radiation. In general, it is difficult to predict in which tissues, or at which life stages, gene expression variation gives rise to observed adaptive differences. We examined transcription in the liver, an easily accessible, metabolically active, tissue. The liver expresses many genes with potential roles in the physiological adaptation to different aquatic environments, including hormone receptors and genes involved in osmoregulation, energy homeostasis, and response to hypoxia. Further, many eQTL identified in this study may be common to other tissues. In general, the extent to which eQTL are shared amongst tissues remains unclear, due to the need for very large sample sizes and the limitations of the statistical methodologies available to address this question (The GTEx Consortium 2015). In particular, variation in gene expression levels amongst tissues means that the power to detect underlying eQTL also varies amongst tissues. Although studies have suggested that up to 70% of genes may have common underlying eQTL across tissues (Nica *et al.* 2011), there is also some evidence that *trans* eQTL hotspots in particular may act in a tissue-specific manner (Grundberg *et al*. 2012). Thus, replication of this study in a greater range of tissues, and at different life stages, would shed more light on the regulatory genetic architecture underlying the parallel changes observed when marine sticklebacks independently colonize freshwater.

To investigate the potential genetic mechanisms generating the nine observed eQTL hotspots we searched for associated loci with known transcriptional regulatory functions, and performed upstream regulator analysis for the genes with eQTL in the hotspots. Although the pathways regulating transcription are still poorly characterized for most genes, particularly in non-mammalian species, these analyses can provide useful preliminary information. We found no evidence that eQTL hotspots were due to the presence of a single ‘master’ regulatory locus, or a cluster of regulatory genes, at the hotspot locations. Although many genes with roles in transcriptional regulation were present in, or regulated by, hotspots, finding such genes is not unexpected: approximately 18% of the human orthologues of BROAD stickleback genes are annotated with the GO terms that we used to identify transcriptional regulators. It is also possible that the regulatory elements generating such hotspots are not annotated coding genes: microRNAs and long non-coding RNAs are potentially important *trans* regulators (Vance and Ponting 2014) and not yet well characterized across the stickleback genome.

Our results suggest that, alternatively, these hotspots may be generated by a complex interaction of multiple transcription regulators. Several well-characterized regulatory proteins were identified as important upstream regulators for genes with *trans* eQTL mapping to the hotspots. Unsurprisingly, these included three genes - HNF4A, ONECUT1 and HNF1A - known to be master transcriptional regulators in the mammalian liver (Odom *et al*. 2004). HNF4A and ONECUT1 were identified as a particularly strongly enriched upstream regulators when examining all genes with a *trans* eQTL at genome-wide p<0.021 (Table S7), and were also found to be enriched when examining the subsets of genes with *trans* eQTL mapping to the hotspots on Chromosome 4, Chromosome 6, and Chromosome 12 (Table S7). None of the three genes were physically located in any hotspot, and we were unable to identify significant eQTL underlying variation in their expression (ONECUT1 was not on the microarray). However, we note that HNF4A is less than 300 kb from hotspot Chr12b. These regulators likely act through direct and indirect interactions with other proteins to regulate transcription. Interacting molecules that are especially of interest in respect to hotspot locations are hypoxia inducible factor 1α and catenin beta-1 (HIF1A & CTNNB1, *trans* regulated by the Chr6 hotspot, Fig. 2), histone deacetylase 3 (HDAC3, located in the Chr4b hotspot, Fig. S4), and vitamin D receptor (VDR, located in the Chr12c hotspot, Fig. S4).

The protein HIF1A has previously been investigated as a selective target of local adaptation in fish. It is part of a transcriptional complex (HIF) that alters the expression of numerous genes in many tissues in response to low oxygen conditions (Nikinmaa and Rees 2005, Liu *et al*. 2013). It is also involved in temperature adaptation in fish (Rissanen *et al.* 2006; Liu *et al.* 2013). Thus, HIF1A is of relevance when fish colonize aquatic environments with differing oxygen regimens, for example benthic vs. limnetic habitats or different areas of the Baltic Sea. Rytkönen *et al.* (2007) found no association between variation in the HIF1A coding region and adaptation to hypoxic conditions across various fish species, and markers linked to HIF1A do not appear be under directional selection amongst Baltic Sea stickleback populations (Shimada *et al.* 2011); however the gene was recently found to be under positive selection in high-altitude loach lineages (Wang *et al.* 2015). HIF1A is known to be transcriptionally regulated in fish (Liu *et al.* 2013), and our identification of a *trans* eQTL for HIF1A demonstrates that regulatory variation for this gene is present in Baltic Sea sticklebacks and could be an alternative, unexamined, target of selection. The proteins HNF4A, CNNB1 and HDAC3 are also involved in the hypoxia response (Xu *et al.* 2011; Wang *et al.* 2015).

In conclusion, we have performed the first genome-wide characterisation of the regulatory architecture of gene expression in *G. aculeatus.* We found that variation in gene expression was influenced by polymorphism in both *cis*-acting and *trans* acting regulatory regions. Trans-acting eQTLS clustered into hotspots. In general these hotspots did not co-locate with regions of the genome known to be associated with parallel adaptive divergence among marine and freshwater threespine sticklebacks. However, one hotspot overlapped with a known QTL underlying salinity tolerance, a locally adaptive trait. Hotspots locations appeared to be mediated by complex interactions amongst regulator molecules rather than the presence of few ‘master regulators’. Our broad-scale study suggests many avenues for finer-scale investigation of the role of transcriptional regulation in stickleback evolution.

## Data accessibility

Raw and normalized microarray data, in addition to R scripts describing the normalization procedure, are available in the ArrayExpress database (www.ebi.ac.uk/arrayexpress) under accession number E-MTAB-3098. RAD sequence reads for each individual have been deposited in the NCBI Sequence Read Archive under BioProject ID PRJNA340327. Input files and scripts will be deposited in DRYAD.

## Acknowledgements

This work was funded by grants from Academy of Finland to C.P., E.L., M. N. and J.M. (grant nos. 129662, 134728, 133875, 136464, 141231, 250435 and 265211). Tiina Sävilammi wrote the script for barcode splitting and we thank her for bioinformatics advice. We also thank the numerous people who helped in obtaining and maintaining sticklebacks. Generous computing resources were provided by the Finnish Centre for Scientific Computing (CSC-IT). Research was conducted under an ethical license from the University of Helsinki (HY121-06). Comments from two anonymous reviewers greatly improved the manuscript.

